# A TRANSLATIONAL LC-MS/MS FRAMEWORK FOR LIPID BIOMARKER IDENTIFICATION AND QUANTIFICATION IN HUMAN PLASMA

**DOI:** 10.64898/2026.04.16.718601

**Authors:** Mark David, Klaus-Peter Adam, Desmond Li, Xin Ying Lim, John G. R. Hurrell, Simon Preston, David A. Peake, Amani Batarseh

## Abstract

Lipid metabolism is increasingly recognized as a hallmark of cancer, yet translating lipidomic discoveries into clinically actionable biomarkers remains constrained by analytical variability and limited standardized validation frameworks. This challenge is further compounded by a chicken-or-egg problem, where expensive standards and labelled internal standards are required to identify and quantitate target lipids, but the diagnostic importance of these targets is uncertain until they can be reliably measured. Previous work had indicated the potential of 48 lipid biomarker species for the prediction of breast cancer from plasma samples using high resolution liquid chromatography mass spectrometry. This study aimed to identify each of these 48 species and develop a quantitative method to determine the absolute concentrations of these lipids in plasma to provide the basis for the development of a clinical assay for use in breast cancer detection. In doing so, we present a pragmatic workflow that bridges lipid discovery with lipid identification and robust quantitative analysis. A curated library of 48 lipid species was established using authentic standards to verify plasma lipids through retention-time matching and high-resolution spectral comparison. In plasma, 41 lipids were confidently identified based on co-elution with standards and diagnostic fragment ions. Method qualification, including assessment of accuracy, precision, recovery, and linearity, was performed across all 48 lipids in parallel with identification, and 46 lipids ultimately met all predefined qualification criteria. Notably, practical constraints, including time, cost, and availability of authentic standards, necessitated performing identification and targeted method development in parallel, highlighting challenges inherent to translating lipidomics into commercial or clinical assays. This workflow provides a reproducible framework for harmonizing lipid identification and quantification, enabling the reliable integration of lipidomic data into biomarker discovery and clinical applications.

## INTRODUCTION

Lipids serve many essential metabolic functions in our bodies, including energy storage, formation of cellular and subcellular membranes, and cell signaling^1^. Aberrant lipid metabolism is increasingly recognized as a hallmark of cancer development and progression across many tumor types including breast^2–10^, colorectal^11–16^, pancreatic^17–20^, lung^21–23^ , prostate^9,24^ and ovarian^25,26^. In breast cancer, altered lipid profiles have been identified both in tumor tissue and in circulating blood. These lipid changes not only mirror the altered metabolism of cancer cells but also open opportunities to develop lipid-based diagnostic and surgical tools. One such innovation is the MasSpec Pen (MSPen) device which leverages differences in lipid composition between tumor and normal breast tissue to enable more precise intraoperative identification of tumor margins during breast-conserving surgery^27^. Beyond the tumor itself, characteristic changes in circulating lipid profiles have also been observed, distinguishing individuals with breast cancer from those without. These systemic lipid alterations hold potential as minimally invasive biomarkers for early detection and disease monitoring. Despite the growing body of evidence of lipid dysregulation in breast cancer, translation of these findings into clinical diagnostic practice has been limited, in part, by methodological and practical challenges. Accurate assessment of participant exposures, such as medication use^28–31^ , supplement intake^32–34^, diet and lifestyle factors^35,36^ can affect lipid profiles in population studies. Furthermore, pre-analytical variables including participant condition at collection, choice of blood tube type, sample handling conditions (duration and temperature), and monitoring for hemolysis can all influence the stability and reproducibility of lipid metabolite measurements ^37–44^. Well planned clinical studies must be designed and executed to control such factors thus ensuring reliable and biologically meaningful results.

The clinical translation of lipidomics-based assays has been hampered by significant analytical challenges. Although liquid chromatography (LC) coupled with high-resolution mass spectrometry (HRMS) is now routinely used for metabolite and lipid discovery, the diversity of analytical strategies, the structural and chemical complexity of the lipidome, and the wide dynamic range of lipid concentrations in biological matrices complicate extraction, separation, and detection^45^. Lipid species vary from highly hydrophilic to highly hydrophobic, making simultaneous analysis difficult and often requiring compromises in chromatographic or ionization conditions. High-resolution mass spectrometry (HRMS)-based workflows can employ several complementary acquisition strategies, including parallel reaction monitoring (PRM), and data-dependent acquisition (DDA). In both PRM and DDA modes, specific precursor (MS¹) ions detected in the survey scan are isolated and fragmented to generate tandem mass spectra (MS²) for structural elucidation and confident lipid identification. When coupled with liquid chromatography LC, these approaches collectively constitute LC-HRMS/MS analyses, which integrate chromatographic separation with high-resolution tandem mass spectrometry. DDA with an inclusion list represents a discovery-driven LC-HRMS/MS approach, in which fragmentation is triggered for pre-defined targets and other targets only when they rank among the most intense ions in the survey scan, while PRM is a fully targeted technique that isolates and fragments specific, pre-defined precursors regardless of their intensity^46–50^. Although these complementary strategies can greatly enhance confidence in lipid identification, achieving complete structural resolution across diverse lipid classes remains an ongoing challenge^51,52^.

Robust quantification for biomarker discovery therefore depends on validated assays that combine accurate lipid annotation, calibration against authentic standards, and reproducible performance across studies^53^. While quality assurance and quality control (QA/QC) guidelines are well established in metabolomics, their consistent application in lipidomics is still emerging, and community-driven initiatives are ongoing to define best practices for human plasma analysis^54,55^. Resources such as the LIPID MAPS consortium provide standardized lipid nomenclature^56^, while dedicated software tools (e.g., LipidSearch^57,58^, Thermo Scientific) enable automated annotation of large and complex high-resolution LC-MS^n^ datasets^59,60^ and support relative quantification using stable isotope-labeled (SIL) internal standards (ISTDs). Together, these significant analytical challenges underscore the need for rigorous, standardized frameworks to generate reproducible and reliable lipid measurements for biomarker discovery and translation into clinical assays.

In our previous work, breast cancer-associated plasma lipid biomarkers were discovered and annotated using untargeted LC-HRMS analysis^61^. In this present work, we aim to verify these 48 lipid biomarkers annotated in pooled human plasma using authentic reference standards and HRMS spectral matching. Building on these verified lipid targets, we then developed a targeted quantitative liquid chromatography tandem mass spectrometry (LC-MS/MS) assay employing lipid reference compounds and SIL ISTDs to generate multi-point calibration curves for each analyte. Overall, the work provides a proof-of-concept framework that integrates rigorous lipid identification with quantitative assay qualification of lipid biomarkers, bridging discovery lipidomics and large-scale clinical translation.

## METHODS

### Materials

#### Reagents and Solvents

Butylated hydroxy toluene (BHT), acetonitrile (ACN), methanol (MeOH), 2-propanol (IPA), water, and formic acid Optima™ LC/MS grade, were purchased from Thermo Fisher (Waltham, MA, USA). Acetone, ammonium formate LiChropur™ (AmF), dichloromethane (DCM), ethanol (EtOH), ethyl acetate (EtOAc) and methyl-*tert*-butyl ether (MTBE) were purchased from Sigma-Aldrich (St. Louis, MO, USA). Medronic acid was purchased from Merck (Darmstadt, Germany) and lithium chloride (LiCl) from Tokyo Chemical Industry Co., Ltd (Portland, OR, USA).

#### Analytical Standards

A total of 48 reference compounds and 18 SIL ISTDs were obtained from commercial suppliers (ANSTO, Avanti Polar Lipids, Cayman Chemical, Echelon Biosciences, Sigma-Aldrich, and Toronto Research Chemicals), with CER d18:1/24:0-d₄ and LPC 14:0-d_4_ synthesized by Precion Inc. (Morrisville, NC, USA). All SIL-ISTDs used were labelled with deuterium. Standard stock solutions were dissolved in 1:1 v/v DCM-IPA (1.00 mg/mL) unless otherwise stated: Lysophosphatidic Acid (LPA) standards in 9:1 v/v IPA-water, and Sphingosine-1-phosphate (S1P) in 4:4:1 v/v/v (DCM-IPA)-MeOH-water, both with sonication. The 48 target lipids and their corresponding SIL-ISTDs are listed in Supplemental Table S1. A combined intermediate (INT) mixture of standards was prepared by combining 100 µL of 48 standard stock solutions and diluting to 10 mL with EtOH (10.0 µg/mL each). Eleven calibration spiking solutions (CSS) were prepared by serial dilution of the INT mixture in EtOH to generate the calibration levels ranging from 0.02-10 µg/mL. SIL-ISTD stock solutions (1.00 or 0.10 mg/mL) were combined and diluted in EtOH to prepare a working ISTD (WIS) solution (1.00-10.0 µg/mL).

#### Columns for reversed phase chromatography

Zorbax Eclipse Plus C18 column (1.8 μm, 2.1 × 100 mm; Agilent, Santa Clara, CA, USA) for reference standard acquisition and lipid identification studies. Combination of C18 1.8 μm columns, 2.1×100 mm and 2.1x50 mm, for method validation.

#### Biological samples

Quality control (QC) samples were prepared from a lot of pooled K_2_EDTA human plasma purchased from BioIVT. Aliquots of 400 µL were stored in 1.5 mL Eppendorf tubes at -80°C until use.

#### Two-phase Lipid Extraction

Two-phase lipid extraction was performed as outlined in Table 1. Plasma (20 µL) was combined with equal volumes of working internal standard solution (WIS) and EtOH. Calibration standards were prepared by replacing plasma with water, followed by addition of WIS and CSS. Blanks contained water and ethanol, with ISTD blanks additionally containing WIS. Protein precipitation and lipid extraction were carried according to a modified MTBE extraction protocol proposed by Matyash and co-workers^62^. Following addition of 50 µL of acidified MeOH-formic acid-BHT (100:1:0.01, v/v/w), samples were vortexed for 1 min, then extracted with 200 µL of water and 400 µL of MTBE. After vortexing for 5 min and centrifugation at 4000 rpm for 10 min, the upper organic layer (250 µL) was collected and evaporated to dryness. Samples were reconstituted in 100 µL of 5:3:1 v/v/v IPA-acetone-(MeOH-formic acid-BHT, 100:1.0:0.01) prior to LC-MS/MS analysis.

**Table 1.**
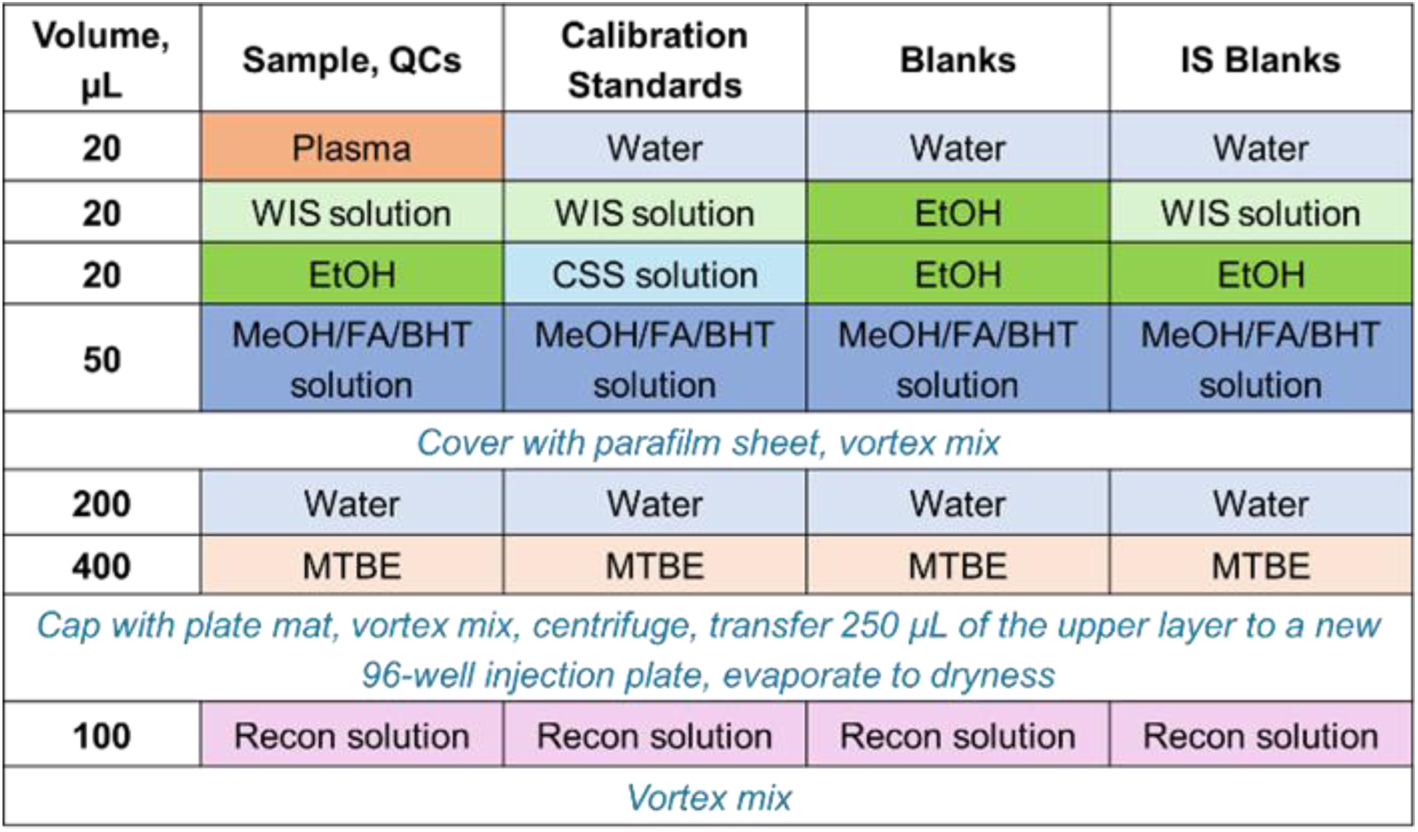
Sample preparation workflow. Lipid extraction was performed on samples, QC samples, calibration standards, blanks, and ISTD blanks using a modified Matyash MTBE protocol. Plasma (20 µL), WIS, and acidified MeOH/formic acid/BHT were combined, followed by liquid-liquid extraction with water and MTBE. The upper organic phase (250 µL) was evaporated and reconstituted in IPA/acetone/MeOH prior to LC-MS/MS analysis.

#### Library generation

Individual lipid standard solutions (5 µg/mL) were analyzed separately by LC-HRMS/MS (Method 1 and Method 2) in both positive and negative ionization modes. Chromatographic mobile phases were supplemented with modifiers, AmF, formic acid, and LiCl, to promote adduct ion formation under electrospray conditions. Instrumental parameters were optimized to allow detection of common protonated, ammoniated, sodiated, and lithiated species in positive mode, as well as deprotonated, chloride, and formate adducts in negative mode. The same conditions facilitated the formation of in-source fragment ions and lithium-associated cluster ions that were evaluated as potential diagnostic species for lipid class confirmation.

This data was analyzed using TraceFinder v5.2 SP1 (Thermo Scientific, San Jose, CA) software by targeting reference compounds by accurate mass, isotope pattern and retention time. Then, the lipid mass spectra were examined manually using FreeStyle software v1.8 SP2 (Thermo Scientific, San Jose, CA) to confirm elemental composition and assignment of negative and positive adduct ions. Finally, lipid adduct ions yielding high-quality MS² spectra were used to generate corresponding mass spectral library entries in mzVault v2.3 SP1 (Thermo Scientific, San Jose, CA).

#### Lipid identification in plasma

Reference standard mixtures (RS1-RS5) were prepared by serial dilution of 50 µg/mL stock solutions in 5:3:1 (v/v/v) IPA-acetone-(MeOH containing formic acid and BHT, 100:1.0:0.01), yielding final concentrations of 0.008–5.00 µg/mL per lipid. For lipid identification experiments, pooled human plasma from previously analysed cohorts was reconstituted in 100 µL of the same solvent system. Non-spiked plasma extracts (P) contained no reference standards or working internal standard (WIS) solution, whereas P0 contained only WIS, and extracts P1-P5 were spiked with both WIS and RS mixtures to match the concentrations of RS1-RS5.

Lipid identification in plasma was based on combined chromatographic and mass-spectrometric criteria to ensure high-confidence structural assignment. Identification required:

i. detection of the molecular (precursor) ion in plasma above the established signal-to-noise threshold;
ii. an MS² fragmentation spectrum matching the corresponding reference standard in the mzVault library, including diagnostic fragment ions and neutral losses;
iii. retention-time matching the authentic standard within the accepted tolerance; and
iv. co-elution of the reference standard mixture with the plasma extract, producing a corresponding increase in intensity at the same retention time, along with a high-quality MS² spectral match and MS¹ mass accuracy within ±2 ppm of the theoretical or reference standard value.

Automated data processing was performed in TraceFinder to integrate LC-HRMS chromatograms, confirm elemental composition and isotopic pattern, and perform preliminary spectral-library matching. All putative identifications were manually reviewed in mzVault, and retention time and MS² spectra were verified in FreeStyle software against authentic reference-standard data.

#### LC- (HR-MS/MS and MS/MS) Analysis

Reference standards and pooled plasma used for the lipid identification study were analysed by LC-HRMS using a Vanquish Horizon Duo ultra-high-pressure liquid chromatography (UHPLC) system (Thermo Scientific, San Jose, CA, USA) coupled to an Orbitrap Exploris 240 high-resolution mass spectrometer (Thermo Scientific, San Jose, CA, USA) with two acquisition methods, DDA (with inclusion list of lipids from standard database) and PRM. A final targeted LC-MS/MS method for lipid quantification and assay qualification was performed using a SCIEX Exion UHPLC system coupled to a 7500 triple quadrupole mass spectrometer (SCIEX, Ontario, Canada).

#### Chromatographic Conditions

All analyses employed reversed-phase separation on Zorbax Eclipse Plus C18 columns maintained at 60 °C. The autosampler compartment was held at 10 °C, and the injection volume was 5 µL. Although all separations utilized reversed-phase conditions, four distinct LC methods were applied depending on the analytical objective (Supplemental Tables S2A, S2B, S3A and S3B). Methods 1 and 2 were used for high-resolution (HRMS/MS) library generation and lipid identification studies, while Methods 3 and 4 were used for the targeted LC-MS/MS assay.

##### Method 1 (Library generation and lipid identification)

Mobile phase (MP) A consisted of 5:3:2 (v/v/v) water–ACN–IPA and contained 0.1% formic acid, 400 mg/L AmF, 1 mg/mL LiCl, and 1 mg/L medronic acid. Mobile phase (MP) B consisted of 8:1:1 (v/v/v) IPA–ACN–ethyl acetate (EtOAc) and contained 0.1% formic acid, 400 mg/L AmF, and 1 mg/mL LiCl.

##### Method 2 (PC/PE isomer identification)

MP A was 5:3:2 (v/v/v) water–ACN–IPA and contained 0.1% formic acid, 400 mg/L AmF, and 1 mg/L medronic acid. MP B was 9:1 (v/v) IPA–ACN and contained 0.1% formic acid and 400 mg/L AmF. For compatibility, EtOAc and LiCl were omitted from MP B, and a longer equilibration period at 0.4 mL/min was applied.

##### Method 3 (Targeted assay qualification, positive and negative ion modes)

Chromatographic conditions were equivalent to Method 1, with a 10 min gradient for the acquisition of positive ion LC-MS/MS data covering acyl carnitine (CAR), lysophosphatidylcholine (LPC), lysophosphatidylethanolamine (LPE), PC, PE, and S1P lipid species, and negative ion data for lysophosphatidic acid (LPA), LPC, lysophosphatidylinositol (LPI), PC, phosphatidylinositol (PI), and phosphatidylserine (PS) lipid species.

##### Method 4 (Targeted assay qualification, targeted positive ion mode)

Also based on Method 1, this 4 min gradient was used for the acquisition of positive-ion LC-MS/MS data for CER, sphingomyelin (SM) and triacylglyceride (TG) species.

#### Mass Spectrometry

##### HRMS/MS (DDA with inclusion list)

Reference standards and plasma extracts were analyzed in both positive and negative ion modes on the Orbitrap Exploris 240 mass spectrometer. Full-scan MS¹ spectra were acquired at 120,000 resolution (FWHM at *m/z* 400) with internal mass calibration active, followed by data-dependent MS² scans at 30,000 resolution using stepped higher-energy collisional dissociation (HCD) with the energies optimized for each ionization mode. Target precursor *m/z* values of reference standard adduct ions were included in a mass inclusion list, with dynamic exclusion set to N = 1 (4 seconds) and a precursor *m/z* filter tolerance of ± 10 ppm.

##### HRMS/MS (PRM)

PRM experiments (analysed on the Orbitrap Exploris 240 mass spectrometer) were performed to determine the retention times and fatty acyl compositions of PC and PE isomers in plasma extracts. Experiments were conducted using UHPLC Method 2 in negative ion mode. Survey MS¹ scans were acquired at 60,000 resolution (FWHM at *m/z* 400), followed by sequential MS² scans at 30,000 resolution using stepped HCD energies. MS² spectra were collected across each lipid’s elution profile to enable full fatty acyl characterization and differentiation of isomeric species.

##### MS/MS (Multiple Reaction Monitoring - MRM)

Two targeted MRM methods were developed on the triple quadrupole SCIEX 7500 system (curtain gas 40 psi, CAD gas 11). Both methods employed a target cycle time of 1000 ms, pause time of 5 ms, and settling time of 15 ms, with maximum dwell times of 250 ms. Method 3 (9.6 min duration; minimum dwell 2 ms) operated in both positive (5500 V) and negative (4500 V) modes at 550°C; Method 4 (4.0 min duration; minimum dwell 3 ms) used positive mode only (5500 V) at 75°C. The MRM settings and transitions for all target lipids and ISTD are summarised in Supplemental Tables S4A - S4C.

##### Targeted assay qualification

Assay qualification metrics are summarized in Table 2, including intra- and inter-assay accuracy, precision, recovery, and stability, evaluated using three batches of QC samples analyzed on different days. The calibration standards were prepared and analyzed across three independent runs conducted on separate days. Depending on the analyte, up to 11 calibration levels were included, each analyzed in duplicate, with a minimum of six levels required to meet calibration acceptance criteria. Each batch also included 12 QC plasma samples. Relative recovery was assessed using six QC controls and six QC samples spiked with middle CCS. Carryover was evaluated by injecting an internal standard blank immediately after the highest calibration level. Benchtop (BT) and freeze/thaw (FT) stability were further assessed using pooled QC plasma samples. The control condition, which served as the reference for both stability assessments, comprised samples that had undergone a single FT cycle, were thawed on ice, and extracted immediately. For BT stability, separate aliquots were (i) thawed on ice and held for an additional 3 h before extraction, (ii) thawed at ambient temperature for 3 h, or (iii) thawed at ambient temperature for 6 h, with timing initiated at the start of thawing. For FT stability, samples were subjected to up to three additional FT cycles, each involving complete thawing on ice followed by refreezing at −80 °C.

**Table 2.** Summary of target lipids identified in plasma. Lipid identifications in plasma were confirmed by matching the reference standard retention time and MS^2^ library spectrum. Lipids that were not confirmed were either not detected due to low abundance in the plasma extract, lacked an MS^2^ library match or did not match the reference standard retention time.

| Lipids confirmed in plasma |  |  |  | Lipids not confirmed |
| --- | --- | --- | --- | --- |
| CAR 18:2 | LPE 18:0 | PE 16:0_18:1 | SM d18:1/18:1 | LPA 16:0 |
| CER d18:1/18:0 | LPE 22:6 | PE 16:0_20:4 | SM d18:1/24:0 | LPA 18:0 |
| CER d18:1/24:0 | PC 14:0_16:0 | PE 18:0_18:2 | S1P d18:1 | LPI 18:1 |
| LPA 18:2 | PC 16:0_16:1 | PE O-18:0_20:4 | TG 16:0_18:1_23:0 | PG 18:0_18:1 |
| LPC 14:0 | PC 16:1_16:1 | PE P-16:0_18:2 | TG 18:1_18:1_18:2 | PI 18:2_20:4 |
| LPC 16:0 | PC 18:0_18:0 | PI 16:0-18:1 | TG 18:1_18:1_22:0 | PS 18:0_18:1 |
| LPC 18:0 | PC 18:0_18:2 | PI 18:0_18:1 | TG 18:1_18:1_22:1 | SM d18:1/17:0 |
| LPC 18:2 | PC 18:1_18:1 | PI 18:0_20:4 | TG 18:2_18:2_18:1 |  |
| LPC 18:0 | PC 18:2_18:2 | PS 18:0_20:4 | TG 18:2_18:2_18:2 |  |
| LPC 18:2 | PC 18:1_20:4 | SM d18:1/18:0 | TG O-16:0_18:1_18:2 |  |
| LPC O-16:0 |  |  |  |  |

## RESULTS

### Overview of analytical workflow

The overall analytical workflow linking lipid discovery, lipid identification, and targeted qualification is outlined in Figure 1. In our previous study, breast cancer-associated plasma lipid biomarkers were discovered and annotated using untargeted LC-HRMS/MS analysis. In the present work, this framework is extended through the generation of a lipid reference library, enabling confident identification of candidate lipids, alongside targeted qualification using LC-MS/MS. This stepwise approach provides a traceable pathway from discovery-based annotation to quantitatively qualified measurement, ensuring that the selected lipid biomarkers are analytically robust and suitable for subsequent clinical translation.

**Figure 1.**
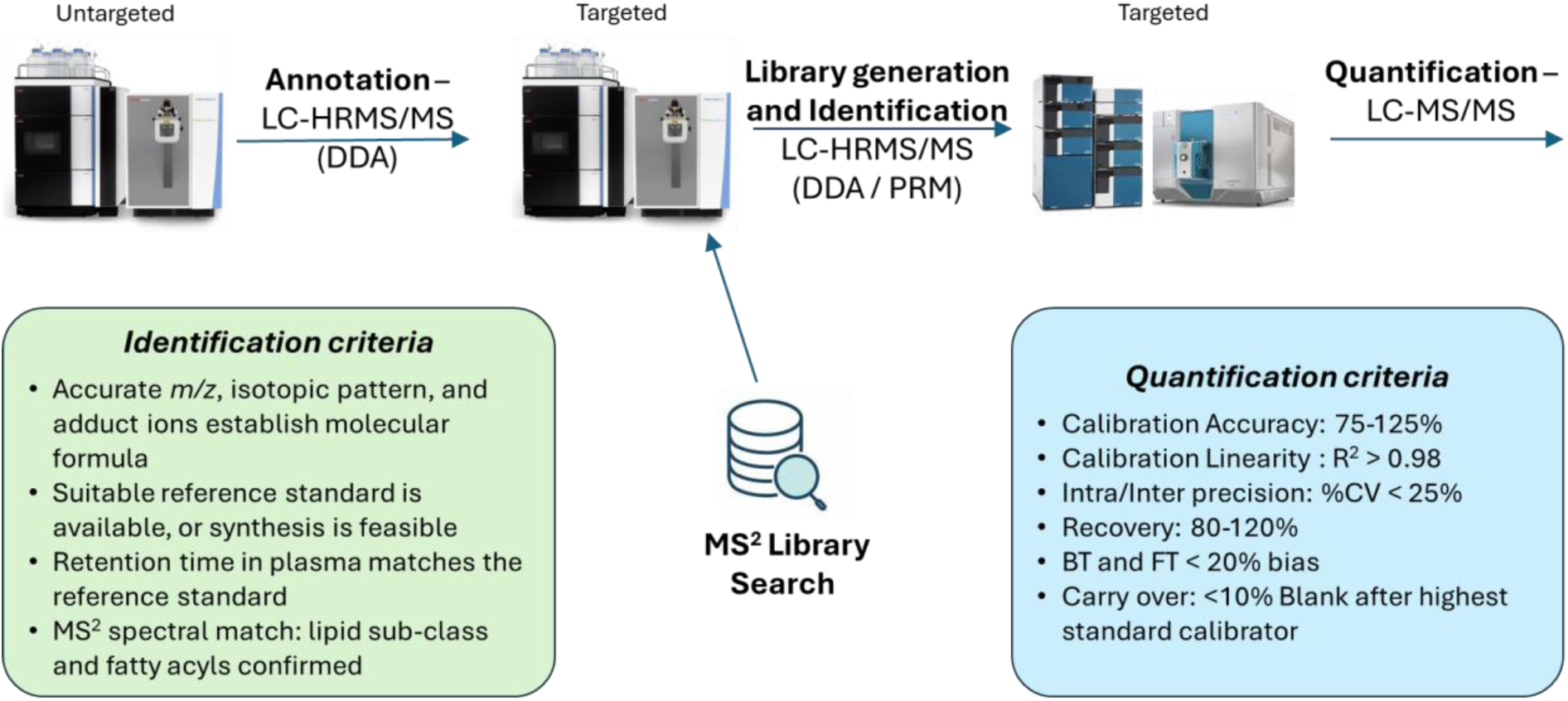
Overview of a framework for lipid biomarker selection, identification and quantification. Untargeted LC-HRMS/MS was used to discover and annotate plasma lipids that differentiate breast cancer vs. controls. Lipid targets were selected from commercially available or synthesized reference standards, and a high-resolution library was created. Lipid identification was performed by LC-HRMS/MS (DDA) using the criteria shown on the lower left, while LC-HRMS/MS (PRM) was applied to confirm the presence of isomeric species. Finally, a quantitative LC-MS/MS method was developed using a triple quadrupole mass spectrometer and qualified using the criteria shown on the lower right.

### Library Generation and lipid identification in plasma

There were a total of 48 reference compound and 18 SIL internal standard database entries used in this study, which comprised 322 MS^2^ spectra.

Analysis of individual lipid standards confirmed the formation of characteristic adduct and fragment ions under the optimized conditions. In positive ion mode, [M+H]⁺, [M+Li]⁺, [M+NH₄]⁺, and [M+Na]⁺ species were detected, together with [M+H-H₂O]⁺ and [M+H-FA]⁺ fragments. In negative ion mode, [M-H]⁻, [M+Cl]⁻, and [M+HCOO]⁻ adducts were observed, along with neutral-loss fragments (M-CH₃) from formate adducts of phosphatidylcholines. Lithium cluster ions, including [M-2H+Li]⁺, [M-H+Li+Cl]⁻, and [M-H+Li+HCOO]⁻, were also detected for selected lipid classes.

Manual verification of mzVault library matches, supported by spectral and retention time confirmation in FreeStyle, demonstrated consistent coelution and diagnostic MS² fragmentation for all confirmed lipids. Based on these criteria, 41 of the 48 lipids from the library were confidently identified in plasma (Table 2). These identifications showed excellent agreement with reference standards (Δ retention time < 0.05 min) and high MS² match scores, with precursor ion mass accuracy typically within ±2 ppm, highlighting the analytical precision of the measurements. The lipids LPA 16:0, LPA 18:0, LPI 18:1, PG 18:0_18:1, and PS 18:0_18:1 were not detected, whereas PI 18:2_20:4 and SM d18:1/17:0 did not meet the identification criteria. Refer to Supplemental Table S5 for full lipid identification details.

The lipid derived metabolite linoleoyl-L-carnitine (CAR 18:2) is presented here as an illustration of the identification process.

#### 1. CAR 18:2 mass spectral library entry creation

Both the protonated [M+H]^+^ (*m/z* 424.3421) and lithiated [M+Li]^+^ (*m/z* 430.3503) adduct ions of CAR 18:2 (Figure 2A) were detected with accurate masses and isotopic distributions consistent with the elemental composition C₂₅H₄₅NO₄ (Figure 2B). After re-evaluating the MS² spectra of CAR 18:2 across the [M+H]^+^ [M+Li]^+^ and [M+CHOO]^-^ adduct forms (including the formate adduct, which was analysed but not illustrated due to low abundance) the resulting MS² spectra were incorporated into the mass spectral reference library (Figure 2C). The same approach was applied to all other lipid species included in the library, using the most informative adduct forms and characteristic fragment ions for each lipid sub-class.

**Figure 2.**
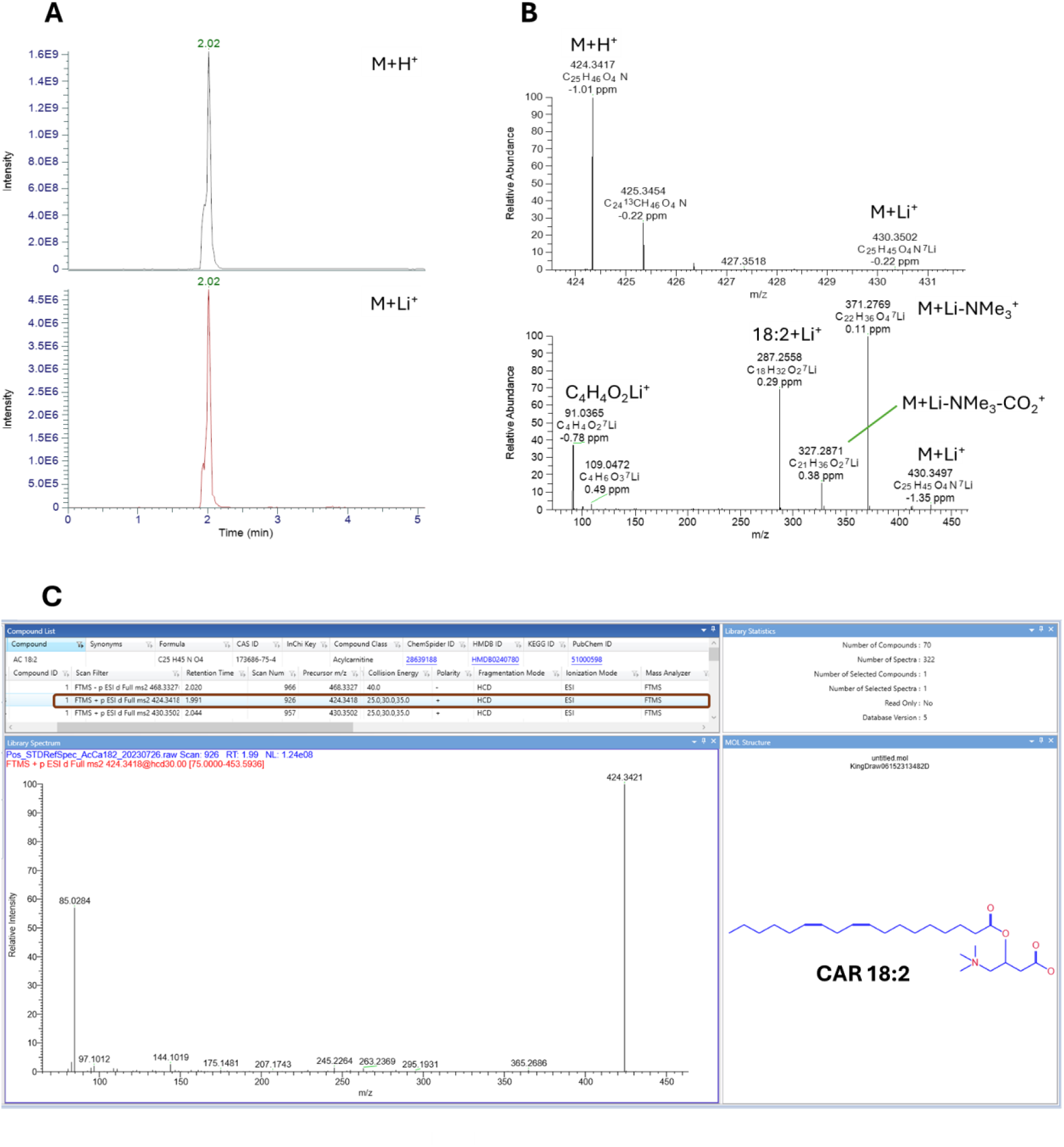
**A**. LC-HRMS/MS chromatograms of CAR 18:2 [M+H]+ *m/z* 424.3421 and [M+Li]+ *m/z* 430.3503. **B**. Mass spectrum of CAR 18:2 and MS² spectrum of [M+Li]+ *m/z* 430.3503. **C**. MS² library entry for CAR 18:2 [M+H]+ *m/z* 424.3421. Protonated and lithiated ions at 2.02 min (**A**) and their isotopic patterns (**B**) confirm the elemental composition, while the MS² of the lithium adduct confirms the 18:2 acyl chain and carnitine headgroup. The library entry (**C**) includes MS² spectra of the protonated, lithiated, and formate adducts, showing the characteristic *m/z* 85.0284 fragment, neutral losses of trimethylamine (*m/z* 365.2686) and fatty acid (*m/z* 144.1019), and the fatty acyl ion at *m/z* 263.2369. Peak broadening in (**A**) is due to high concentration and signal saturation.

#### 2. Identification of CAR 18:2 in plasma

Automated identification of CAR 18:2 in neat plasma using TraceFinder software is illustrated in Figure 3. The results status in the software indicates that CAR 18:2 is identified in the neat plasma sample with [M+H]^+^ *m/z* 424.3422, 0.20 ppm mass error, found at 1.99 min, with a 100% isotopic pattern score matching the composition of C_25_H_45_NO_4_. The MS^2^ spectrum in plasma (Figure 3A), matches the library entry for CAR 18:2 (Figure 3B) with a match score of 90.

**Figure 3.**
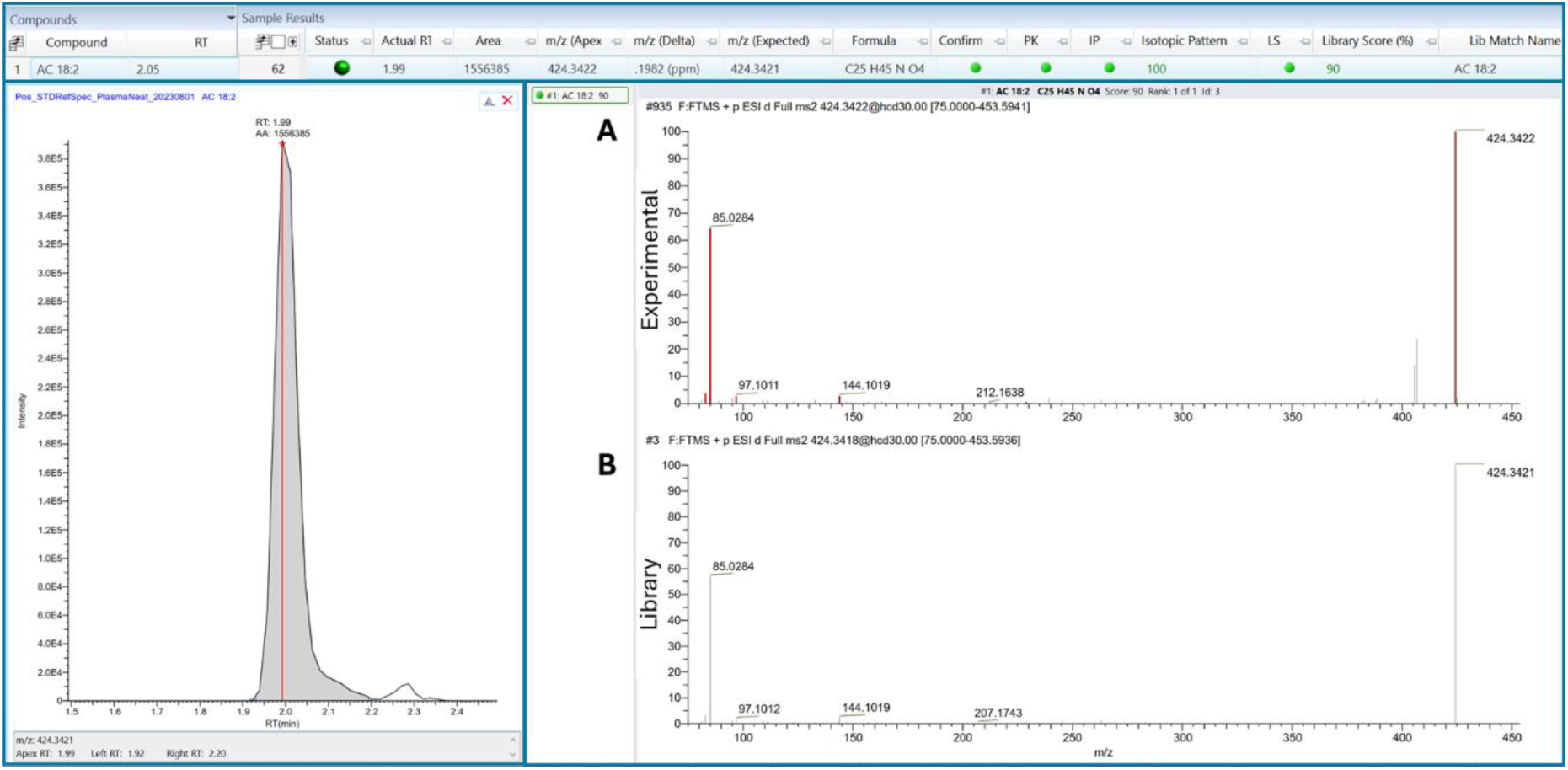
Automated LC-HRMS/MS Identification of CAR 18:2 in human plasma. **A.** MS^2^ spectrum of *m/z* 424.3422 in plasma. **B**. MS^2^ spectrum of CAR 18:2 *m/z* 424.3421 in the mass spectral library. Results displayed in TraceFinder software confirm the peak at 1.99 min. with accurate mass and isotopic pattern matching the elemental composition C_25_H_45_NO_4_ and a positive library match to CAR 18:2 with a score of 90.

The retention times of the CAR 18:2 standard (2.02 min), neat plasma (1.99 min), and plasma spiked with the standard (1.99 min) (Figures 4A, 4A, and 4D, respectively) confirm that the compound identified in plasma coelutes with the reference within ±2 seconds. In addition, the [M+H]^+^ signal intensity increased from 4.0e^5^ in plasma to 1.0e^8^ in the spiked plasma extract. As expected, CAR-d_3_-18:2 (*m/z* 427.3610) also co-elutes with CAR 18:2 (Figure 4D) and the measured *m/z* of d_0_ and d_3_ protonated molecular ions and ^13^C isotopes fit to the expected elemental composition within ±1.0 ppm error.

**Figure 4.**
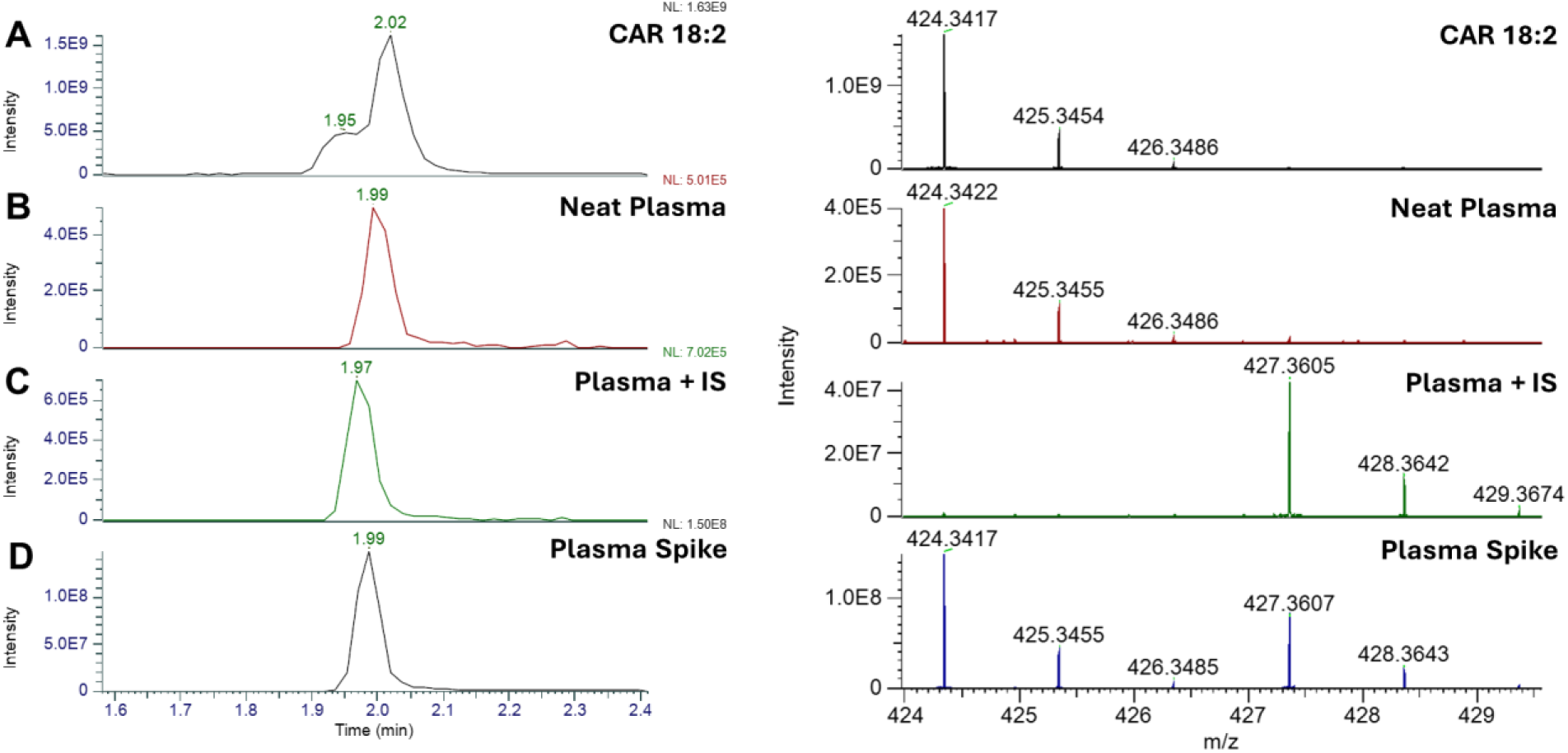
Confirmation of CAR 18:2, *m/z* 424.3421, retention time and mass spectrum in: **A**. CAR 18:2 standard, **B**. neat plasma, **C.** plasma spiked with d_3_-ISTD, *m/z* 427.3610, and **D.** plasma spiked with reference standard RS1. The retention time of the target peak in plasma (1.99 min.) matches the standard (2.02 min.) and spiked plasma sample (1.99 min) within ±0.03 min. As expected, peak intensity increases from 4.0e5 in plasma to 1.0e8 in plasma spiked with CAR 18:2.

### Challenges with Isomeric species

Targeted LC-HRMS/MS (DDA) analysis of plasma lipids revealed that several PC, PE and TG species were present as mixtures of isomers. For example, seven PC and PE lipids comprised two to four isomeric species, as determined by manual inspection of the negative ion MS² spectra. The DDA approach provided broad coverage of lipid features; however, a key limitation of this method is that repeated scans occur primarily on the precursor or MS¹ level. As a result, the MS² data acquired across chromatographic peaks were often insufficient for complete resolution and identification of isomeric diacyl phospholipids. To address this limitation, targeted LC-HRMS/MS analyses were subsequently performed using a PRM approach to identify the isomeric fatty acyl anions associated with each PC and PE lipid group. The PRM experiment for the PC 36:2 species is summarised in Supplemental Table S6. LC-HRMS/MS chromatograms of the [M+HCOO]⁻ adduct ion (*m/z* 830.5917), together with the corresponding MS² spectrum of PC 18:1/18:1 at 7.55 min, are shown in Figure 5A. As expected, the only fragment ion detected was *m/z* 281.2486, corresponding to the 18:1 fatty acyl anion. Similarly, analysis of the PC 18:0/18:2 standard (Figure 5B) yielded fragment ions at *m/z* 279.2329 and 283.2642 at 7.68 min, consistent with 18:2 and 18:0 fatty acyl anions, respectively.

**Figure 5.**
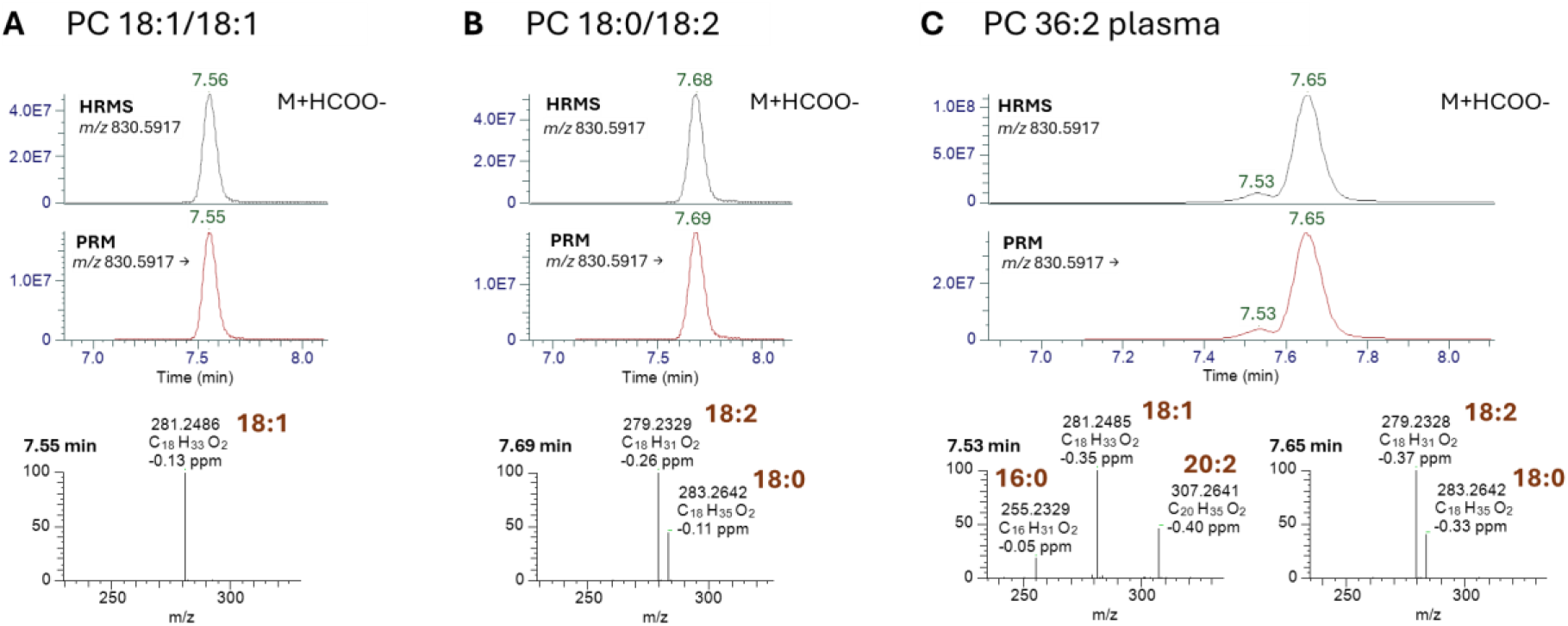
LC-PRM identification of PC 36:2 isomers in human plasma. Targeted LC-HRMS/MS analysis of PCs in plasma and standards. **A**. PC 18:1/18:1 standard showing the [M+HCOO]⁻ adduct and corresponding fragment ion at *m/z* 281.2486. **B**. PC 18:0/18:2 standard with fragment ions at *m/z* 279.2329 and 283.2642, corresponding to 18:2 and 18:0 fatty acyl anions. **C**. Pooled plasma PC 36:2 showing a mixture of isomeric species, with fragment ions at *m/z* 255.2329, 281.2845, and 307.2641 corresponding to PC 18:1_18:1 and PC 16:0_20:2 for the first peak, and *m/z* 279.2328 and 283.2642 corresponding to the abundant peak, PC 18:0_18:2.

In contrast, analysis of pooled plasma revealed that PC 36:2 comprised a mixture of isomers. Fragment ions at *m/z* 255.2329, 281.2845, and 307.2641 at 7.53 min (Figure 5C) corresponded to PC 18:1_18:1 and PC 16:0_20:2 isomeric species. The sn-position and double bond location of the fatty acyl chains could not be determined from the MS² spectra. Additionally, the [M+HCOO]⁻ adduct ion at *m/z* 852.5760 (7.65 min) produced ions consistent with PC 18:0_18:2. The minor 0.02-0.03 min retention time shift relative to the reference standard is within the expected experimental variation for partially coeluting chromatographic peaks that were not fully resolved by the separation method.

### Targeted Assay Qualification

Assay qualification metrics are summarised in Table 3. The qualification consisted of four batches of QC sample analyses for determining accuracy and precision of calibration standards, precision, relative recovery, carryover and pre-analytical stability (BT and FT). Qualification criteria and results are briefly described here.

1. **Calibration.** Calibration concentration ranges for the quantified analytes were initially estimated and implemented across a broad concentration range. From this, the linear dynamic range for each lipid was empirically determined to ensure the method was fit for purpose. The resulting calibration performance, including linearity and accuracy, is summarized in Supplemental Tables S7, S8, and S9, respectively. Calibration curves demonstrated good linearity (R² > 0.98) for 46 of the 48 lipids. PS 18:0/18:1 and PS 18:0/20:4 were the two exceptions, failing to meet standard calibration criteria (R² < 0.98 and accuracy outside the 75 to 125% window).
2. **Precision**. Intra-assay precision (CV < 20%) was achieved for all analytes except SM d18:1/24:0, TG 18:1_18:1_22:0, and TG 18:2_18:2_18:2, each of which showed a single intra-day measurement slightly outside the acceptance criteria; these values can be considered isolated outliers and do not materially affect overall precision. Inter-assay precision met all acceptance criteria (Supplemental Table S10).
3. **Stability.** All temperature- and time-stressed QC samples met acceptance criteria (<20% bias) on ice and at room temperature for all analytes except for TG 16:0_18:1_23:0, TG 18:1_18:1_18:2, TG 18:1_18:1_22:0, and TG 18:1_18:1_22:1. These four triacylglycerols met acceptance criteria only under the control condition. Therefore, they are considered qualified solely when sample handling follows the established stability control procedure, including thawing from the freezer on ice and immediate extraction (Supplemental Table S11).
4. **Recovery.** Relative recovery was generally consistent across all lipids. Recovery was assessed only for lipids with matched internal standards, as measurements without a deuterated analogue could vary substantially due to matrix effects and ionization differences. For these lipids, recoveries were within 80–120% of the spiked QC concentrations. LPC 16:0 and LPC 18:0 showed apparent lower recovery when calculated using the standard approach (Relative Recovery A) due to their high endogenous levels; applying the adjusted calculation (Relative Recovery B) for these lipids provided a more accurate estimate of extraction efficiency. (Supplemental Table S12).
5. **Carry-over.** Analyte area in blanks following the high calibration standard were ≤ 10% of the analyte area of plasma QCs.

**Table 3.** Assay qualification criteria and results summary. Assay qualification metrics including calibration standards (accuracy and precision), precision of QC samples (inter- and intra-assay coefficients of variation, CV), recovery (relative recovery of spiked QCs), carryover (blank area following high concentration standard), BT and FT stability (bias).

| Metric | Experiment | Analysis performed | Acceptance Criteria | Results |
| --- | --- | --- | --- | --- |
| **Calibration standard (CAL STDs): Accuracy and precision** | Analyse 11 CAL STDs in duplicate, minimum of 3 runs over at least 3 days | Determine % accuracy and inter- and intra-assay precision, %CV | 1) <u>Accuracy</u>: Within 75-125% value of lower limit of quantitation (LLOQ), 80-120% of higher limit of quantitation (ULOQ).<br>2) <u>Precision</u>: Inter-assay % CV ≤ 25% at LLOQ, and ≤ 20% at ULOQ.<br>3) <u>Exclusion of Calibration standards</u>: 75% of standards for each analyte must meet accuracy requirements at every calibration concentration.<br>4) <u>CAL function and weighting</u>: Linear fit ($R^2 > 0.98$) with 1/x or 1/x<sup>2</sup> weighting. | 1. Pass, except for: PS 18:0/18:1 and PS 18:0/20:4.<br>2. Pass<br>3. Pass<br>4. Pass, except for: PS 18:0/18:1 and PS 18:0/20:4 |
| **Precision of QC Samples** | Analyse 12 QCs per analytical run for a minimum of 3 runs. | Determine inter- and intra-assay precision, CV. | Intra-assay and inter-assay CVs ≤ 20% | Pass, except for one intra-day precision for: SM d18:1/24:0, TG 18:1_18:1_22:0, TG 18:2_18:2_18:2 |
| **Recovery** | Un-spiked QC: Analyse 6 aliquots of QC plasma<br>Spiked QC: Analyse 6 aliquots of QC plasma spiked with concentration equivalent to the middle CSS | % Relative Recovery $A = 100 \times (\text{Average conc. in spiked QC samples} - \text{average conc. in plasma QC controls}) \div (\text{spiked CSS})$ | Relative Recovery within 80–120% of the spiked concentration. | Pass except for LPC 16:0 and LPC 18:0, where Relative Recovery B used = $100 \times (\text{Average conc. in QC controls} + \text{nominal spiked CSS}) \div (\text{Average conc. in QC controls})$ |
| **Carry-over** | Include blank sample after ULOQ in each run. | Compare analyte area in blanks after ULOQ to the analyte area of QC plasma. | Analyte area in blanks following ULOQ ≤ 10% analyte area of QC plasma. | Pass |
| **Sample Stability Testing** | <u>Benchtop (BT) Stability</u>:<br>QC samples (n = 6) extracted after 3 h on ice, and 3 or 6 h at room temperature; control = average of non-stressed QCs.<br><u>Freeze-Thaw (FT) Stability</u>:<br>QC samples (n = 6) extracted after 3 FT cycles; control = average of non-stressed QCs. | <u>BT Stability</u><br>$\text{Bias} = 100 \times (\text{BT condition} - \text{Control}) \div (\text{Control})$<br><u>FT stability</u><br>$\text{Bias} = 100 \times (\text{FT condition} - \text{Control}) \div (\text{Control})$ | BT and FT stability, Bias ≤ 20%. | <u>BT Stability</u><br>All Pass, except for TG 16:0_18:1_23:0, TG 18:1_18:1_18:2, TG 18:1_18:1_22:0, and TG 18:1_18:1_22:1<br><u>FT stability</u><br>All Pass, except for TG 16:0_18:1_23:0, TG 18:1_18:1_18:2, TG 18:1_18:1_22:0, and TG 18:1_18:1_22:1 |

Of the 48 lipids targeted by the targeted LC-MS/MS method, 46 met all predefined assay qualification criteria.

## DISCUSSION

This work was designed to demonstrate a comprehensive framework for translating lipidomic discoveries into a robust, targeted quantitative assay. Starting from a discovery panel of candidate lipids, a spectral library was established to support confident lipid identification in plasma samples, followed by development of a targeted LC-MS/MS workflow for quantitative analysis of selected lipids with potential clinical relevance. Under typical circumstances, such studies would be performed sequentially: database curation, lipid identification in discovery experiments, and subsequent development of a targeted assay. In this case, 48 lipids were initially included in the database, 41 were confirmed in plasma, and therefore only these 41 would normally be advanced into targeted quantitation.

However, a well-recognized ‘chicken-or-egg’ problem in lipidomics is that extensive sets of authentic reference standards and stable isotope-labelled (SIL) lipids are often not acquired at the stage of lipid identification, primarily due to their high cost and limited availability. While a fully sequential workflow is generally preferred, as described above, this constraint limited the feasibility of such an approach in the present study. Instead, a parallel strategy was adopted, whereby lipid identification and targeted assay development were progressed concurrently.

Species-specific SIL standards were incorporated where available, and lipid assignment was supported through retention time, MS² fragmentation, and other orthogonal analytical parameters. Under this framework, all 48 candidate lipids were included in the LC-MS/MS assay, rather than restricting analysis to only those confidently confirmed by HRMS. Notably, several lipids that were not confidently identified in plasma by HRMS were detectable in the targeted assay due to its higher sensitivity, highlighting that the “discoverable” set based on HRMS alone would have been more limited. In commercial or applied laboratory settings, such adaptations are often necessary: the combined costs of mass spectrometry operation, consumables, and reference standards, alongside their limited availability, can make a strictly sequential identification-to-quantification process impractical. Consequently, workflows must balance analytical rigor with practical and commercial constraints. Initial lipid identification was hindered by low ionization efficiency, co-eluting isomers, and minor retention-time shifts between plasma-derived species and neat standards. These differences are consistent with matrix effects or unresolved isomeric interferences. DDA facilitated structural assignment through characteristic fragment ion spectra, but it could not fully characterize isomeric species. PRM experiments were subsequently performed to obtain diagnostic fragment ions that enabled confident phospholipid isomer discrimination.

Nonetheless, some ambiguity remained, underscoring the need for orthogonal separation techniques or advanced fragmentation strategies in future method iterations. Achieving higher levels of structural assignment can be appropriate at the right stage of assay development. The authors believe that complete molecular characterization should be performed once a clinical assay has been fully established. This can be done using a variety of methods to achieve positional or regio-isomeric characterization, such as as electron-induced dissociation (EID)^63–68^, multistage MSⁿ ^69^, and/or ultraviolet photodissociation (UVPD) ^70–78^. While these approaches offer superior structural resolution, we don’t believe this is required during assay development and at the stages discussed in this paper. In this study, rigorous verification using authentic standards and spike-in experiments was performed, improving confidence in lipid identification and an important step toward clinical implementation. Importantly, the goal of this study was not exhaustive structural elucidation but to demonstrate a feasible and reproducible workflow suitable for eventual clinical use. Within this context, the data demonstrates that the defined lipid panel can be robustly qualified, with measurements carefully controlled using internal standards and calibration to ensure accuracy and consistency. These results confirm that, despite the challenges of translating discovery data into quantitative assays, reliable qualification of a lipid panel is achievable.

The LC-MS/MS method demonstrated robustness across multiple qualification parameters, including linearity, accuracy, precision, recovery, and stability. Acceptance criteria were informed by established bioanalytical validation frameworks, including the U.S. Food and Drug Administration, European Medicines Agency, and Clinical and Laboratory Standards Institute, providing a structured basis for defining reasonable, fit-for-purpose performance expectations and enabling future validation to proceed within recognized regulatory expectations. Most lipids met these qualification criteria, including those quantified using surrogate internal standards from the same lipid class. However, PS 18:0_18:1 and PS 18:0_20:4 did not qualify, likely reflecting the use of non-class matched surrogates, which could not adequately correct for analyte-specific ionization differences. These limitations were compounded by inherently poor ionization efficiency and minor retention-time variability for these species. Overall, these outcomes highlight the practical constraint of representing 48 target lipids with only 17 internal standards, a necessary compromise given the limited commercial availability of deuterated analogues and the time and cost associated with custom synthesis. In some cases, authentic standards also required tailored synthesis for structural verification, further increasing analytical complexity and cost. Collectively, these constraints underscore the challenge of translating lipid discovery into quantitative assays, necessitating a balance between analytical breadth, quantitative rigor, and practical feasibility for real-world implementation.

Future development will require expanding the internal standard panel so that each lipid is paired with its own labelled analogue, with analytical method validation performed under strict regulatory compliance using real clinical samples processed and handled as they would be in routine practice. Combined with the eventual adoption of advanced structural elucidation tools as they become more accessible, these refinements will be key to moving from qualification to full validation. From a broader perspective, this work reflects the evolving landscape of translational lipidomics, where methodological robustness and standardization are increasingly prioritized over exploratory depth. Efforts such as those led by the Lipidomics Standards Initiative remain important for standardizing lipid identification and improving transparency and consistency in reporting. Continued expansion of curated spectral libraries and shared reference resources will further strengthen reproducibility and comparability of lipid data, facilitating integration of lipidomics into clinical and regulatory pipelines.

## CONCLUSIONS

In this study, we developed a targeted LC-MS/MS workflow that advances lipidomics from putative annotation to robust quantitative analysis. Authentic lipid standards were sourced to generate a high-quality reference database, and spike-in experiments were performed to verify lipid identities through retention-time and MS² spectral matching. A total of 48 lipid species were incorporated into the spectral library and included in the targeted assay. In plasma, 41 lipids were confidently identified based on co-elution with authentic standards and high-quality MS² library matches. Method qualification, including assessment of accuracy, precision, recovery, and linearity, was performed across all 48 lipids in parallel with identification, and 46 lipids ultimately met predefined criteria. Notably, practical constraints, including time, cost, and availability of authentic standards, necessitated performing identification and method qualification in parallel, reflecting real-world challenges in translating lipidomics into clinical assays.

Collectively, this work establishes a proof-of-concept framework that demonstrates progression from lipid identification to assay qualification, while highlighting practical considerations for clinical assay development. Although full analytical validation remains to be completed, the current level of method qualification provides sufficient robustness to enable exploratory application to clinical plasma cohorts. Future studies will apply this workflow to breast cancer and control plasma to investigate lipid biomarkers of clinical significance.

## Contributions

A.B, X.Y.L., D.A.P., K.P.A., H.W., Q.Z., A.L., and D.L contributed to the conceptualization, data acquisition, data curation, formal analysis, and methodology. M.D. wrote the final draft and contributed to review and editing. D.A.P, and J.H contributed to supervision, review, and editing.

## Data Availability

Data is contained within the article or Supplementary Material

## Code Availability

No custom code was generated or analyzed in this study

